# Hybrid Scaffolds Decouple Biochemical & Biophysical Regulation of Cell Phenotype

**DOI:** 10.1101/2025.06.12.659085

**Authors:** Xinyuan Song, Samantha C. Mitchell, Abbie N. Smart, Will Hardiman, Daniel V. Bax, Ceri Staley, Catherine Probert, Pamela Collier, Marian Meakin, Alison A. Ritchie, Tania Mendonca, Amanda J. Wright, Victoria James, Anna M. Grabowska, Cathy L. R. Merry, Serena M. Best, Ruth E. Cameron, Jennifer C. Ashworth

**Affiliations:** Cambridge Centre for Medical Materials, University of Cambridge, Cambridge, UK; School of Veterinary Medicine & Science, University of Nottingham, Nottingham, UK; Translational Medical Sciences, School of Medicine, University of Nottingham, Nottingham, UK; Optics and Photonics Research Group, Faculty of Engineering, University of Nottingham, Nottingham, UK; Ex Vivo Cancer Pharmacology Centre, School of Medicine, University of Nottingham, Nottingham, UK

**Author notes:** Co-corresponding authors:* SMB, REC, JCA.

**Keywords:** Hydrogel, collagen scaffold, ice templating, breast cancer, extracellular matrix

## Abstract

The extracellular matrix changes dramatically during the progression of diseases like cancer. These complex, tissue-specific changes are not adequately replicated by most current biomaterial disease models. Here we demonstrate, for the first time, a biomaterial system allowing combined, independent control over stiffness, composition and 3D collagen architecture. We demonstrate successful perfusion of defined hydrogel formulations into ice-templated collagen scaffolds, controlling the composition of these hybrid scaffolds at constant stiffness. The Young’s moduli of these hybrid scaffolds can also be tuned independently of composition via chemical crosslinking. Encapsulation of human dermal fibroblasts reveals that fibroblast morphology depends on hybrid scaffold composition and on viscoelasticity, highlighting the importance of a system that decouples biophysical from biochemical properties. Finally, we demonstrate application of this system to exert combined control over biochemical and biophysical drivers of cell growth and invasion, focussing on breast cancer as proof-of-concept. We show that collagen fibre patterning enhances breast cancer cell proliferation, also directing the invasion of patient-derived breast cancer cells. These hybrid scaffolds are therefore promising new tools for dissecting the diverse but complementary roles played by the extracellular matrix in regulating cell phenotype, in a range of healthcare applications.

## Introduction

In recent years, increasing recognition of the complex roles played by the extracellular matrix (ECM) in health and disease has led to ever more sophisticated 3D cell culture systems capable of replicating these properties. Many of these systems are based on hydrogel designs of increasing complexity, with several recent studies examining their potential to offer independent regulation over ECM composition and stiffness.^1,2^ Hydrogels are widely used as *in vitro* models of the ECM, given their ability to mimic many properties of natural tissues, including their high degree of hydration and fibrillar architecture.^3^ However, there are inherent limitations. Hydrogels commonly have lower stiffness than that of many soft tissues, and often undergo cell-induced contraction during culture, making quantitative analysis of cell morphology challenging.^4^ Furthermore, in some dense hydrogels, the diffusion rates of nutrients and waste can be limited, inhibiting cell growth or even leading to cell death in the central regions.^5^ Moreover, while techniques for influencing fibre architecture in hydrogels are constantly emerging, currently the microstructural cues presented to cells cannot fully replicate the complex fibre architectures found in living tissues.^6,7^

Ice-templated collagen scaffolds offer an alternative system for 3D culture, with an extensive body of literature exploring their tissue engineering applications.^8,9^ Some key studies also demonstrate their application for disease modelling, particularly for fibrotic diseases such as breast cancer, in which collagen fibre architecture is extensively remodelled.^10,11^ Ice-templating is a proven method for controlling collagen fibre architecture, with the potential to mimic pore sizes and shapes characteristic of tissues ranging from lung to tendon.^12^ Recent advances now allow pore and fibre morphology to be controlled within ice-templated scaffolds more than ever before.^13,14^ However, these too have their own limitations, including relatively low control over ECM composition, and the fact that their sponge-like structure often means that encapsulated cells adhere to single collagen fibres and so are not fully surrounded by ECM ligands in 3D.

In this paper, we demonstrate the controlled combination of defined hydrogels with ice-templated collagen scaffolds, to produce hybrid scaffolds combining the advantages of both systems. We demonstrate application of these hybrid scaffolds to exert independent control over ECM stiffness and composition, while simultaneously tuning collagen fibre organisation. By combining these hybrid scaffolds with patient-derived breast cancer cells, we further validate their application for investigating the role of collagen fibre organisation on breast cancer invasion, in the presence of defined ECM. In this way, we show the suitability of this highly tuneable method for decoupling the diverse biochemical and biophysical ECM influences in both health and disease.

## 2. Results

### 2.1 Hybrid Scaffolds: Combined Control over Biophysical & Biochemical Cues

The design principle behind ice-templated hybrid scaffolds allows independent control over multiple ECM properties in a single system (Figure 1a). Hydrogel perfusion into an ice-templated scaffold of defined fibre structure allows simultaneous control over biophysical properties (stiffness and fibre organisation) and biochemical properties (ECM composition). Here we demonstrate successful hydrogel perfusion into ice-templated scaffolds derived from fibrillar collagen I, focussing on (i) synthetic self-assembling peptide gels (PG) and (ii) collagen gels (CG).

**Figure 1:**
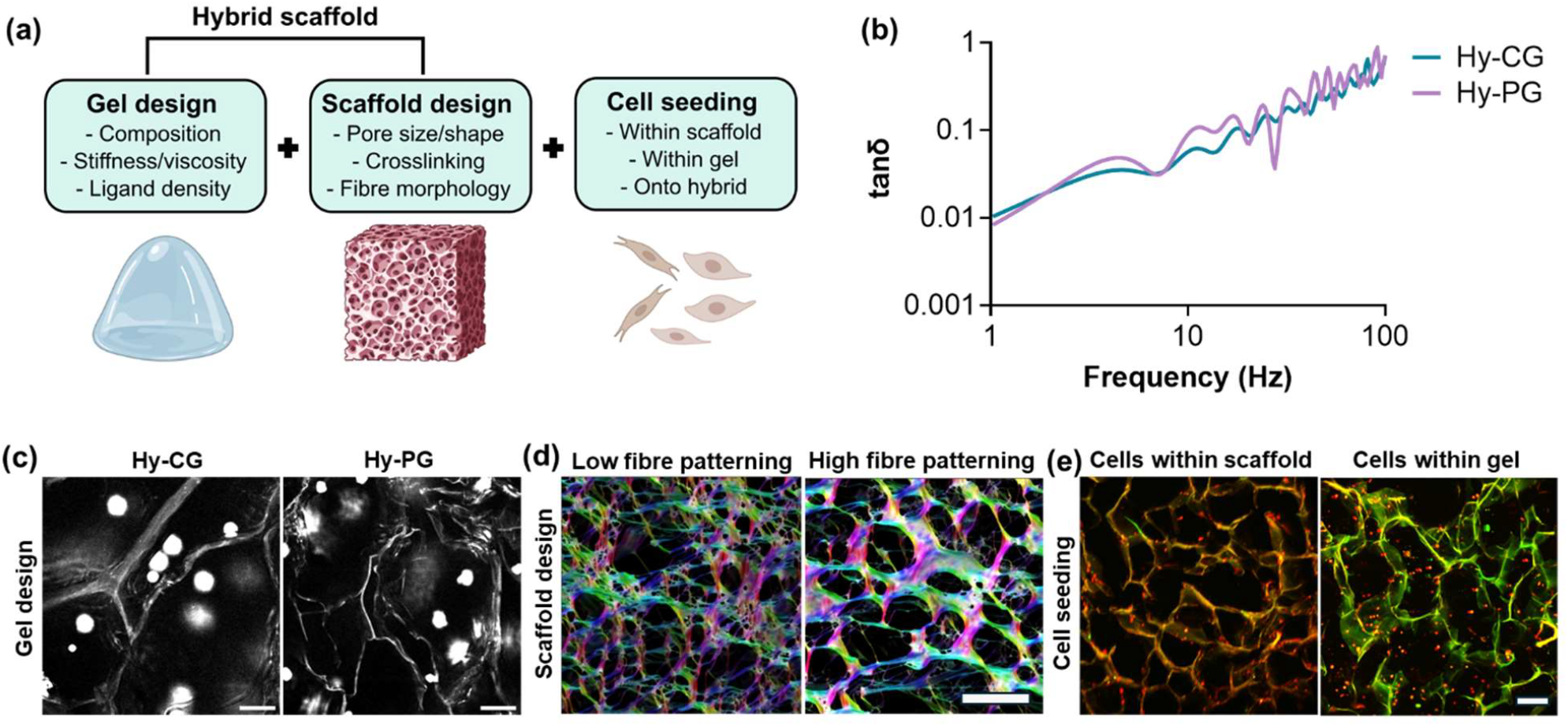
Hybrid biomaterials allow combined control over multiple biophysical properties. (a) Schematic demonstrating degrees of freedom in hybrid scaffold design, incorporating control over collagen scaffold design, gel design, and localisation of seeded cells; (b) Microrheology analysis showing plots of tanδ characteristic of viscoelastic solids for cell-free hybrid scaffolds containing collagen gel (Hy-CG) or self-assembling peptide gels (Hy-PG); (c) Multiphoton images showing efficient penetration of both collagen and self-assembling peptide gels to produce a hybrid scaffold as shown by encapsulated polystyrene beads, scale bar 50 µm; (d) Scanning electron microscopy images with false colour overlay (colour = orientation, brightness = coherency) showing collagen fibre structural control in scaffold design via change of solute during ice-templating (low fibre patterning = 0.001 M HCl, high fibre patterning = 0.05 M acetic acid), scale 50 µm; (e) Multiphoton images of human dermal fibroblasts in hybrid scaffolds, demonstrating that cells may be successfully seeded into the scaffold or the gel component of the hybrid scaffold, scale bar 200 µm.

We found that while hybrid scaffolds containing peptide gels (Hy-PG) could be fabricated using standard gelation protocols,^15^ an amended gelation protocol was required to fabricate hybrid scaffolds containing collagen gels (Hy-CG). Both gel formulations require pH adjustment to induce self-assembly, from either an alkaline (PG) or acidic (CG) precursor solution. Since ice-templated collagen scaffolds incorporate a dilute acid, we found that an amendment to the gelation protocol was necessary for successful Hy-CG fabrication, with additional NaOH required to induce CG neutralisation compared to typical published protocols (Supplementary Figure 1a,b).

Particle tracking microrheology (Figure 1b) demonstrates that PG and CG both undergo self-assembly and gelation within hybrid scaffolds, giving plots of tanδ against frequency typical of viscoelastic solids, where tanδ represents the ratio of the viscous and elastic components in the mechanical response.^16^ Encapsulation of fluorescent beads within the hydrogel component, followed by multiphoton imaging, revealed that these fluorescent beads are evenly distributed throughout the ice-templated collagen structure (Figure 1c, Supplementary Figure 1c), indicating successful hydrogel infiltration and hybrid scaffold formation for both gels tested. Combined with control over the degree of fibre patterning via ice-templating (Figure 1d) and control over the method of cell seeding into the hybrid scaffold (Figure 1e), these hybrid scaffolds therefore represent a highly tuneable platform for 3D cell culture with controlled biophysical and biochemical properties.

### 2.2 Hybrid Scaffolds Provide Independent Control over Stiffness and Composition

To investigate the potential of hybrid scaffold technology for controlling the mechanical environment in 3D cell culture, bulk compression testing was applied to measure the Young’s modulus of each hybrid scaffold, relative to its hydrogel and scaffold components alone. We observed that for both Hy-PG and Hy-CG, the Young’s modulus of the hybrid scaffolds was significantly greater than that of either the hydrogel alone (p<0.01, Figure 2ai) or the ice-templated scaffold alone (p<0.05, Figure 2aii). However, there was no significant difference in Young’s modulus between Hy-PG and Hy-CG, indicating that hybrid scaffold composition can be controllably altered while maintaining a constant compressive stiffness (Figure 2aiii).

**Figure 2:**
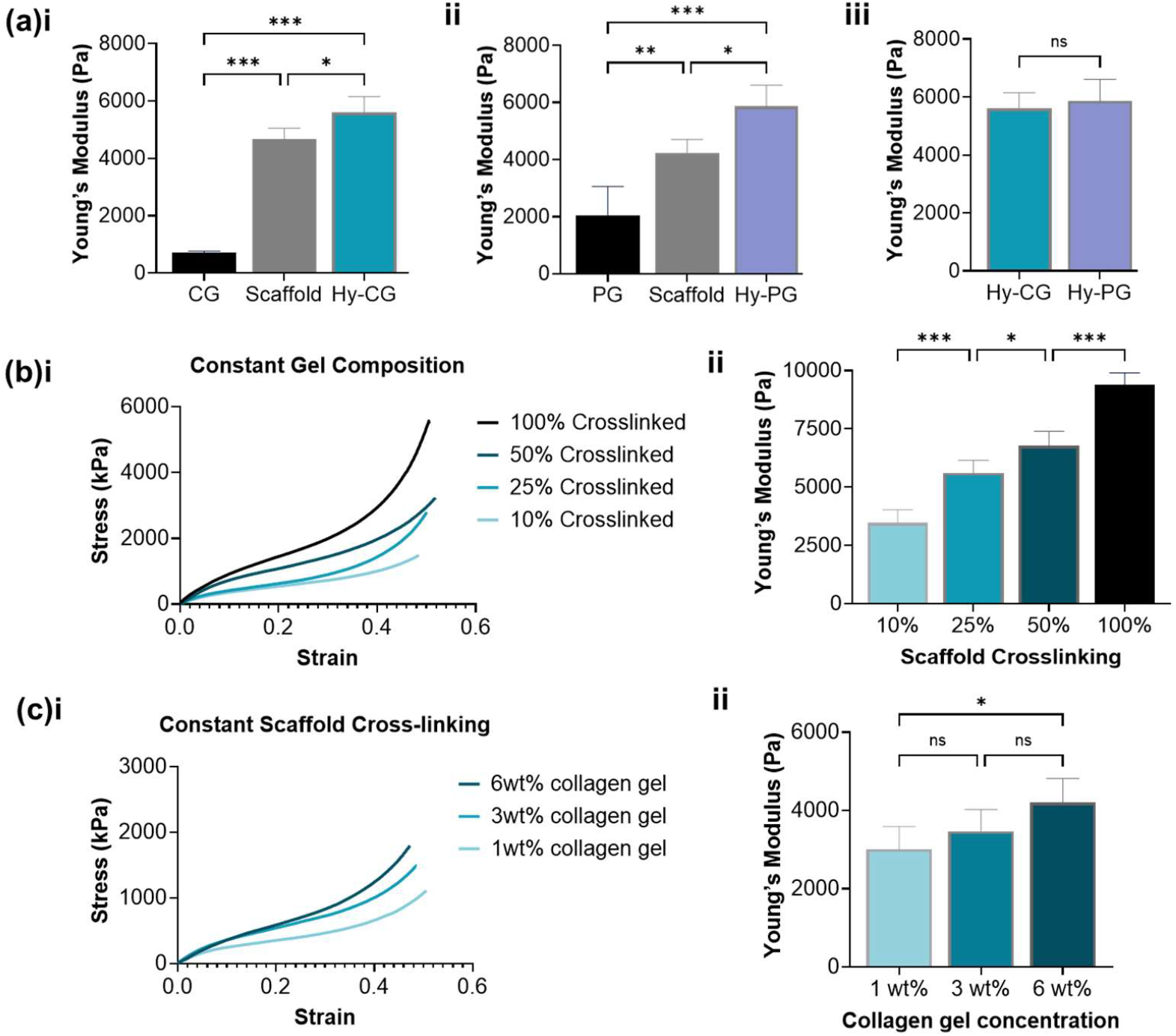
Bulk mechanical properties can be controlled independently of composition. (a) Hybrid scaffolds containing (i) collagen gel (Hy-CG) or (ii) peptide gel (Hy-PG) have significantly higher Young’s modulus than either the gel or scaffold alone, with (iii) no significant difference in Young’s modulus between the two hybrid scaffold compositions; (b)(i) the scaffold component of the hybrid scaffold (Hy-CG) may be crosslinked to different degrees (100% crosslinking refers to the standard 5:2:1 carbodiimide molar ratio) at constant gel composition, producing (ii) significant increases in Young’s modulus; (c)(i) the concentration of the collagen gel component may be controlled, which may produce (ii) modest but significant changes in Young’s modulus, depending on the magnitude of the change in concentration. Error bars represent standard deviation of at least n=4 independent measurements per condition. * p<0.05, **p<0.01, ***p<0.001.

We also investigated control over stiffness in hybrid scaffolds of constant composition (Hy-CG). Tuning the level of carbodiimide crosslinking in the ice-templated scaffolds, prior to hybrid scaffold fabrication, produced controlled changes in Hy-CG mechanical stiffness ranging from 3-10 kPa, with all changes found to be statistically significant (p<0.05, Figure 2b). Finally, we examined control over collagen density in hybrid scaffolds with constant cross-linking (Figure 2c). Increasing the collagen concentration in the CG component produced modest changes in Hy-CG stiffness, which only became significant when collagen concentration was increased by a factor of 6. This therefore indicates a substantial range over which hybrid scaffolds allow independent control over bulk mechanical stiffness and composition, through altering collagen density at constant stiffness, or vice versa.

### 2.3 Fibroblast Morphology is Dictated by Hybrid Scaffold Design

We next examined the influence of hybrid scaffold design on the phenotype of encapsulated cells. Human dermal fibroblasts (HDF), known to elongate within and contract pure collagen gels, were encapsulated within CG alone, scaffolds alone, and Hy-CG hybrids. Strikingly, when HDF were incorporated within Hy-CG, no contraction was seen over culture, in contrast to HDF seeded into CG alone (Figure 3a). This was observed consistently across different concentrations of collagen gel, as well as different degrees of scaffold stiffness as controlled by carbodiimide cross-linking (Supplementary Figure 2a), indicating that these hybrid scaffolds do not undergo macroscale remodelling during HDF culture.

**Figure 3:**
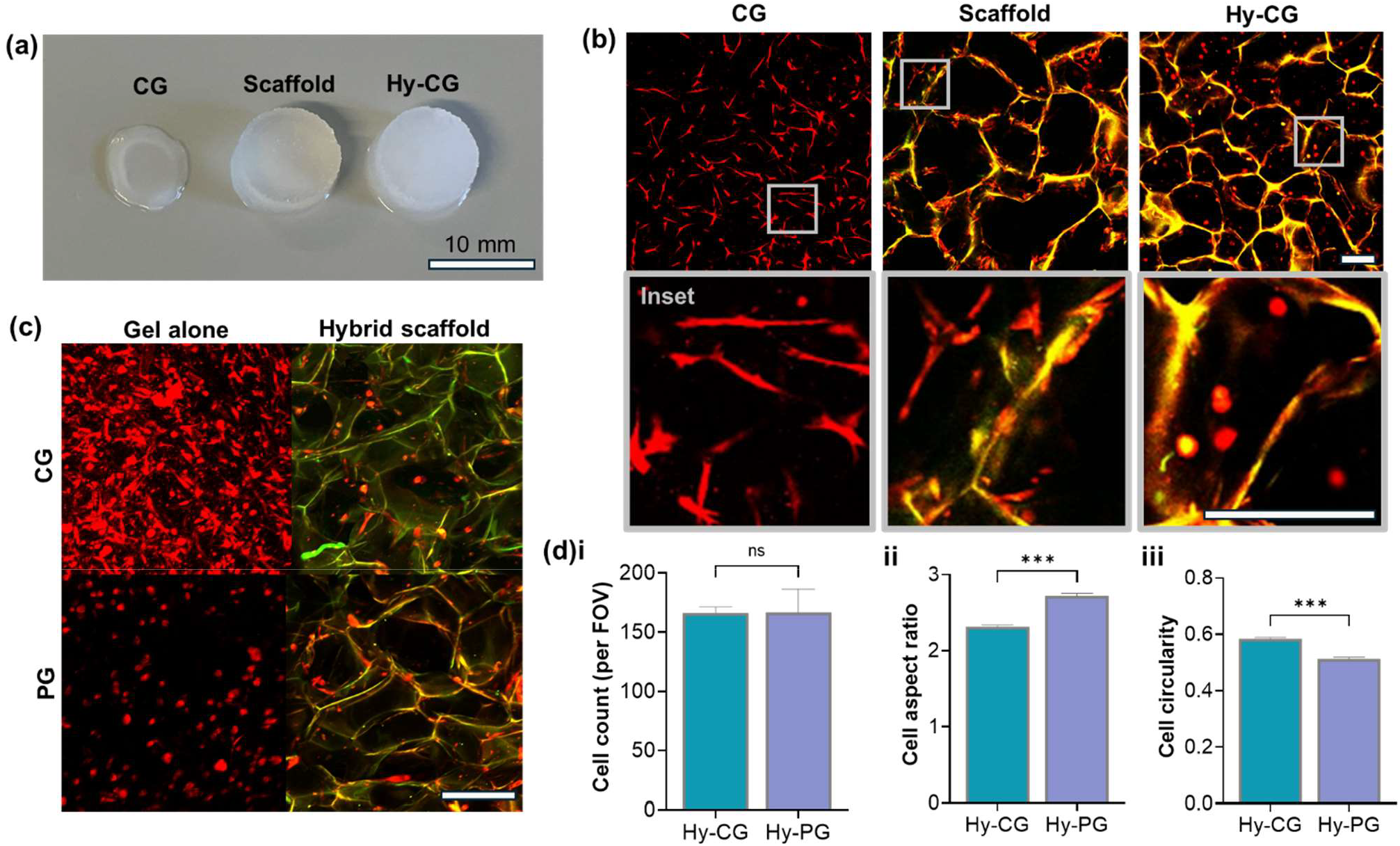
Fibroblast elongation is dependent upon hybrid scaffold composition. (a) Images demonstrating visible contraction of collagen gel (CG) relative to scaffold and hybrid scaffold Hy-CG after 2 days culture with human dermal fibroblasts (HDF); (b) Two photon images of HDF within each material, demonstrating elongation in CG and scaffold alone but a rounded morphology in Hy-CG; (c) Two photon images of HDF after 3 days culture in collagen gel or peptide gel (PG) alone, and corresponding hybrid scaffolds; (d) quantification from n=3 biological replicates shows that (i) cell count is unchanged, (ii) cell aspect ratio is significantly higher in Hy-PG, corresponding to (iii) significantly lower cell circularity, ***p<0.001 All scale bars are 200 µm unless indicated.

Interestingly, multiphoton imaging revealed that, whereas HDF showed a classical elongate morphology in either the CG or scaffold alone, they remained rounded in the corresponding Hy-CG hybrid scaffold (Figure 3b). Again, this result was observed across different levels of scaffold cross-linking as well as at different collagen gel concentrations (Supplementary Figure 2b). The influence of HDF seeding within the scaffold rather than within the CG component of the hybrid scaffold (see Figure 1a schematic) was also investigated. No significant difference was seen in cell aspect ratio or circularity between the two seeding methods, though a significantly higher cell count was observed on HDF seeding within the CG component (Supplementary Figure 3).

In contrast, HDF seeded into Hy-PG had a comparatively elongate morphology (Figure 3c). This result is particularly striking given that HDF exhibit a rounded morphology in the pure peptide gel, consistent with observations in previous studies.^17^ Quantification of HDF morphology showed a significantly lower aspect ratio and higher circularity in Hy-CG relative to Hy-PG (p<0.001, Figure 3d). This indicates that Hy-PG are more conducive to cell-matrix interaction between encapsulated cells and the ice-templated collagen fibres within the hybrid scaffold. However, quantification of cell number after 3 days’ culture revealed no significant difference between Hy-PG and Hy-CG, indicating no difference in ability to support cell proliferation.

To explore this result further, we applied particle tracking microrheology to examine the local mechanical environment within hybrid scaffolds. Micron-sized beads were encapsulated within both CG and Hy-CG (Figure 4a) prior to gelation, and their movement tracked via high framerate video microscopy to measure mechanical properties on the length scale of a single cell. We observed that both the CG and Hy-CG behaved as classical viscoelastic solids, with constant *G’* (storage modulus) and increasing *G’’* (loss modulus) as frequency increases, though the magnitude of *G’’* was consistently lower in Hy-CG relative to CG (Figure 4b). Further analysis of these results demonstrated that while there was a small but non-significant change in *G’*, both *G’’* and tanδ (loss tangent, ratio of *G’’/G’*) were significantly lower in Hy-CG relative to CG (p<0.05, Figure 4c). This indicates that the local mechanical behaviour in the hybrid scaffold is closer to an elastic solid than the pure collagen gel, which has a greater contribution from the viscous mechanical response. Interestingly, no such change was observed in the case of the peptide gel, with no difference in any of *G’, G’*’ or tanδ observed in the comparison between PG and Hy-PG (Supplementary Figure 4).

**Figure 4:**
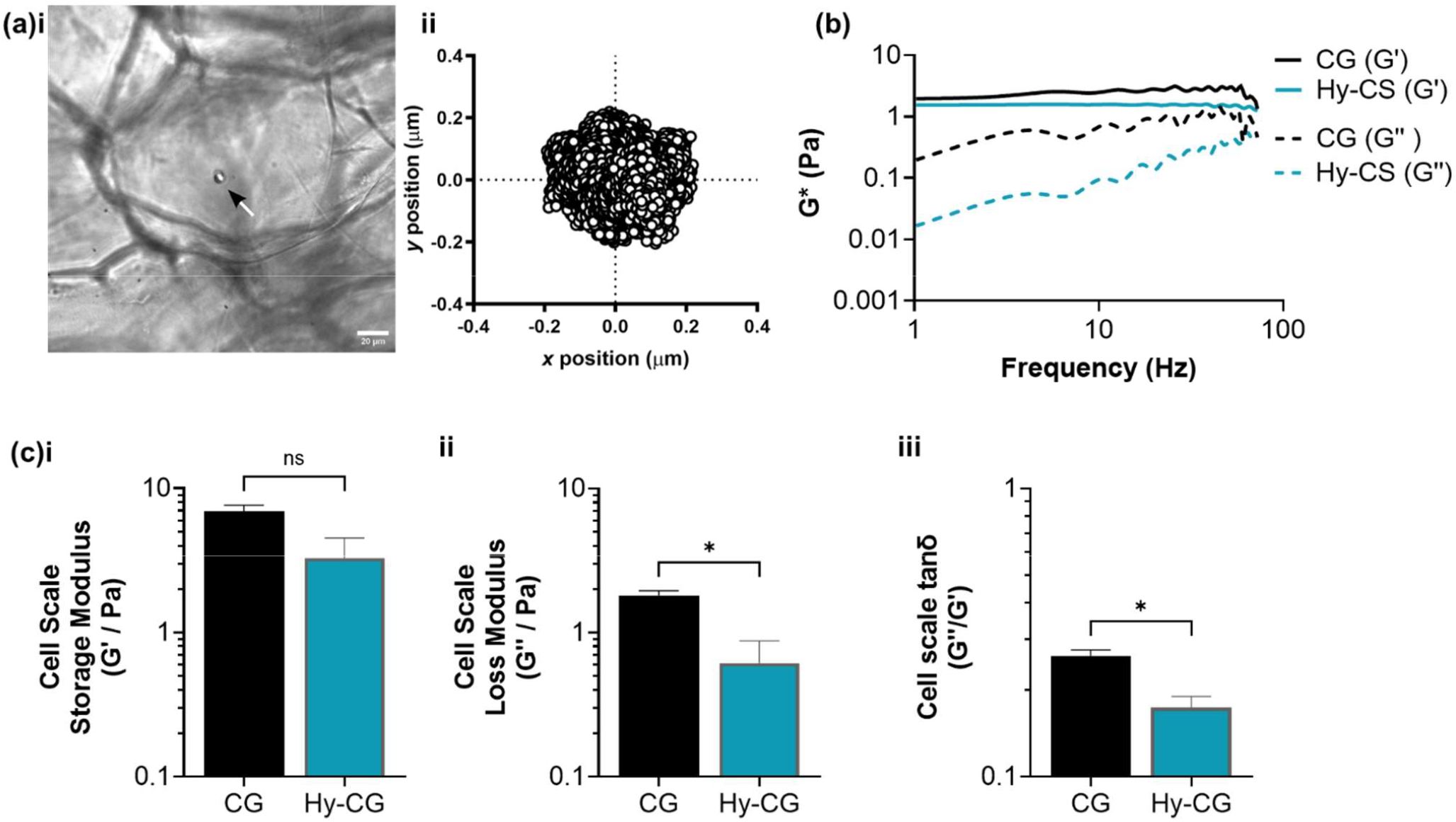
Collagen gels within hybrid scaffolds have altered local viscoelasticity relative to collagen gels alone. (a) Images demonstrating application of particle tracking microrheology in which (i) 6 µm diameter beads (white arrow) are encapsulated within the collagen gel (CG) component of the hybrid scaffold, and (ii) their position tracked via timelapse microscopy; (b) Mathematical analysis of bead trajectory over time gives plots of G* (complex shear modulus) as a function of frequency, revealing a decrease in loss modulus G’’ in the hybrid scaffold; (c) Quantification of (i) G’ (storage modulus), (ii) G’’ (loss modulus) and (iii) tanδ (G’’/G’) reveals a significant decrease in both G’’ and tanδ in the hybrid scaffold relative to the collagen gel alone. Measurements are averaged between 1-100 Hz. Error bars represent standard error of n=3 independent experiments. Scale bar 20 µm.

### 2.4 Hybrid Scaffolds Support and Direct Breast Cancer Cells in 3D Culture

The ability to control biochemical and biophysical properties within hybrid scaffolds indicates their future potential for decoupling ECM influences on cell behaviour across a range of applications. Taking breast cancer modelling as an example application, we investigated the ability of hybrid scaffolds to support breast cancer cell lines in 3D culture. For this investigation, we initially focussed on two well-characterised lines with contrasting morphologies in 3D culture: MCF7, which typically forms round colonies with disorganised nuclei, and MDA MB 231, which display an elongated shape with limited cell-cell interactions.^18^ Hy-PG was chosen for this investigation due to its superior ability to promote cell interaction with the ice-templated collagen fibres (Figure 3), as well as the fact that its viscoelastic response is unchanged relative to PG alone (Supplementary Figure 4).

The PrestoBlue™ viability assay was used to assess cell metabolic activity within the Hy-PG hybrid scaffolds, compared with PG or scaffold alone. While MCF7 was found to proliferate equally well in all conditions tested (Figure 5a), demonstrating a significant increase in metabolic activity over time (p<0.05), MDA MB 231 was more sensitive to the choice of 3D culture environment (Figure 5b). A significant increase in metabolic activity over time was only observed in the scaffold alone or the Hy-PG hybrid, with only a small and non-significant increase in the PG alone. End-point fixation and cytoskeletal Phalloidin staining provided further context to these results, demonstrating a clear distinction between the typical morphology of the two cell lines. In all conditions, MCF7 grew collectively, forming discrete spherical clusters in PG alone, but anisotropic growth following the ice-templated collagen fibres within the Hy-PG hybrid and scaffold alone (Figure 5c). Conversely, MDA MB 231 grew as isolated single cells, which remained rounded in PG alone, but elongated along the patterned collagen fibres within the Hy-PG hybrid and scaffold alone (Figure 5d). This demonstrates the hybrid scaffolds’ ability to direct cell growth and anisotropy, while maintaining a cell type specific morphology.^18^

**Figure 5:**
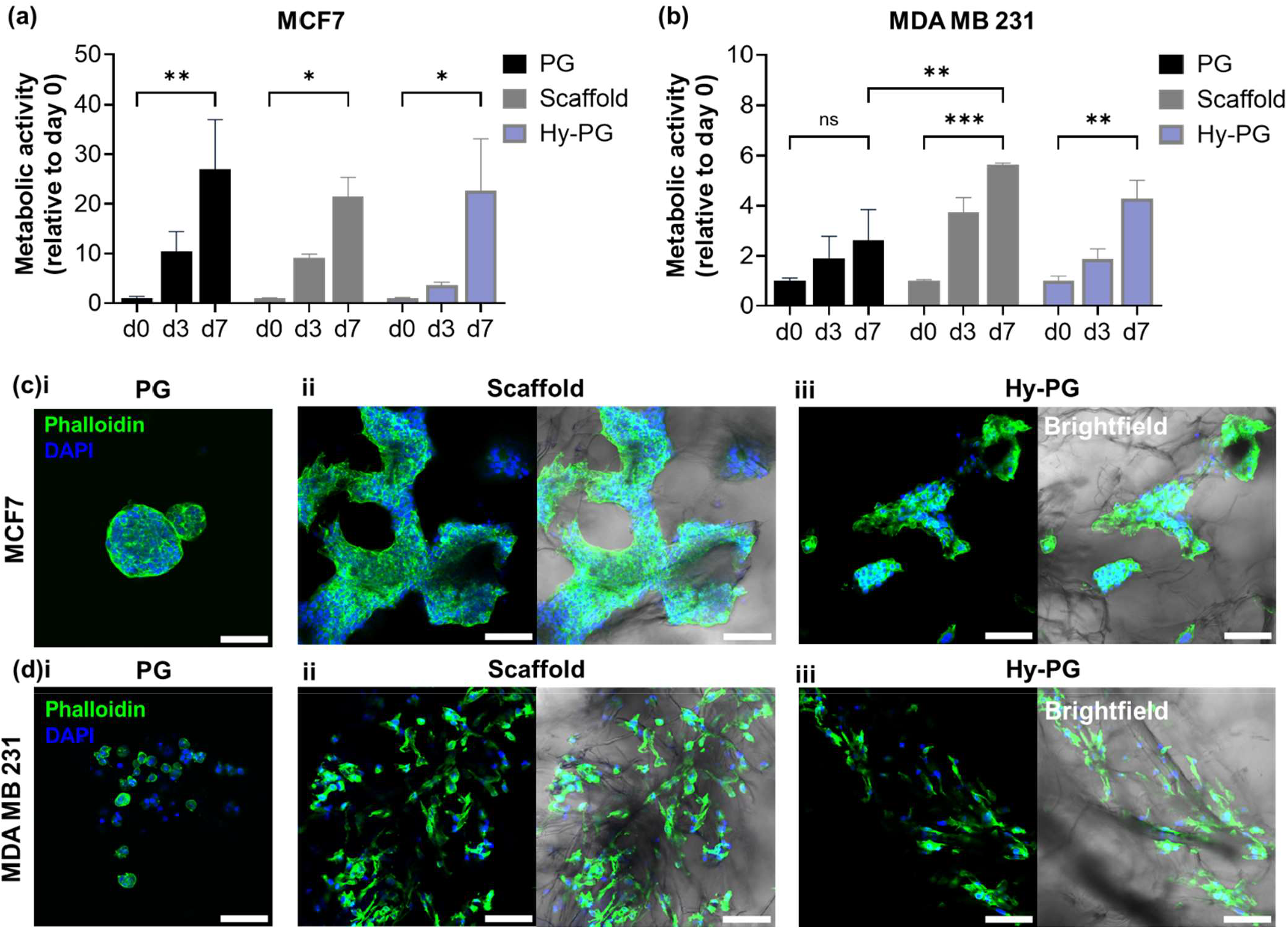
Hybrid scaffolds support breast cancer cell line culture. Presto Blue measurements of metabolic activity for (a) MCF7 and (b) MDA MB 231 encapsulated within peptide gels (PG), scaffold alone, or hybrid scaffolds containing peptide gel (Hy-PG). All conditions support a significant increase in metabolic activity over the culture period, other than MDA MB 231 in PG alone, with significantly lower metabolic activity than in scaffold culture. Fluorescent staining of (c) MCF7 and (d) MDA MB 231 morphology after 7 days of culture reveals that in (i) peptide gel alone, MCF7 grow into discrete clusters while MDA MB 231 remain as single cells with limited cell-matrix interaction. In (ii) scaffold alone or (iii) hybrid scaffolds both cell types show directional growth along the ice-templated collagen pore walls, maintaining cell type specific morphology with MCF7 growing collectively and MDA MB 231 as single cells. Plots show mean and standard error of n=3 independent biological replicates; * indicates p<0.05, ** p<0.01, *** p<0.001. Scale bar 100 µm.

### 2.5 Fibre Structure within Hybrid Scaffolds Directs Primary Breast Cancer Invasion

Incorporation of peptide gels within these hybrid scaffolds opens up exciting potential to tune both ECM composition and structure to direct breast cancer invasion. Applying a previously described approach for controlling ECM composition within defined peptide gels,^17^ we tuned the peptide gel composition to induce local cell invasion of primary breast cancer cells from patient-derived xenografts (PDX, figure 6(a)). This was achieved using additions of human collagen I, recombinant vitronectin and recombinant fibronectin, producing an invasion-permissive peptide gel (iPG). We therefore set out to investigate whether hybrid scaffolds incorporating iPG (Hy-iPG) could form a suitable test system for examining ECM influences on long-range cell invasion.

**Figure 6:**
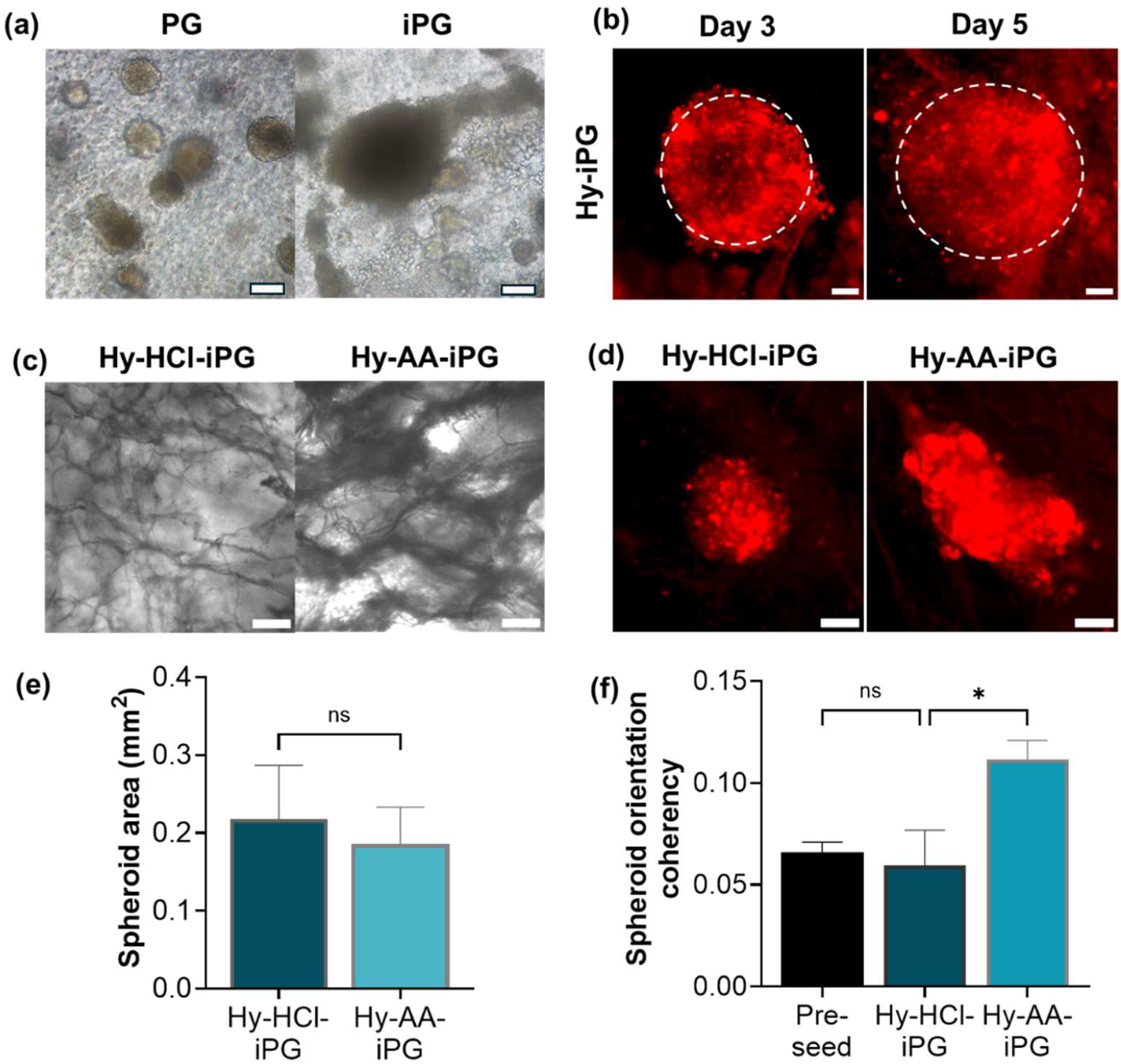
Independent control over hybrid scaffold fibre structure guides primary breast cancer invasion. (a) Peptide gels modified with invasion-promoting ECM additions (iPG) promote increased growth and invasion relative to unmodified peptide gels (PG); (b) Spheroid invasion within hybrid scaffolds incorporating iPG (Hy-iPG) may be tracked over time using CellTrackerTM dye; (c) Fibre morphology in the hybrid scaffolds is controlled via choice of solute during scaffold ice templating (Hy-HCl-iPG: hybrid scaffold incorporating iPG and scaffold templated with HCl; Hy-AA-iPG: hybrid scaffold incorporating iPG and scaffold templated with acetic acid); (d) Control over fibre morphology controls the directionality of spheroid invasion, with Hy-AA-iPG inducing directed invasion along collagen fibres. Quantification of (e) spheroid size and (f) orientation coherency demonstrates significantly higher orientation coherency at constant spheroid size induced by the Hy-AA-iPG hybrid scaffold relative to Hy-HCl-iPG. Plots show mean and standard error of day 5 measurements from n=3 patients; * indicates p<0.05. All scale bars 100 µm.

Spheroids composed of aggregated breast cancer PDX cells were successfully incorporated into hybrid scaffold culture, showing homogeneous growth with maintained spherical morphology over time (Figure 6b). PDX spheroids retained the applied CellTracker™ dye until day 5 (Figure 6b), allowing quantification of spheroid size and morphology over time. We therefore applied these hybrid scaffolds to investigate the role of collagen fibre organisation on PDX invasion. Applying our previously published methods, we varied the solute used for ice-templating of the scaffold component, producing a collagen fibre organisation that either inhibits or promotes cell invasion by the choice of hydrochloric acid (HCl) or acetic acid (AA) as solute respectively.^19^ On incorporation of iPG, this resulted in hybrid scaffolds termed Hy-HCl-iPG and Hy-AA-iPG (Figure 6c). Application of these hybrid scaffolds to PDX invasion revealed that spheroid invasion in the Hy-AA-iPG condition showed higher directionality, corresponding to the presence of collagen fibres, while spheroids in the Hy-HCl-iPG condition grew homogeneously and remained more spherical (Figure 6d). Quantification of spheroid size and orientation coherency demonstrated that while there was no significant efect of fibre structure on spheroid size (Figure 6e), the directionality of invasion in Hy-AA-iPG was significantly greater than in Hy-HCl-iPG (p<0.05, Figure 6f). Spheroids in Hy-AA-iPG showed almost twice the orientation coherency of those in Hy-HCl-iPG, which showed no change in orientation coherency relative to control measurements taken before seeding onto the hybrid scaffolds. This result demonstrates proof-of-concept for application of hybrid scaffolds to reveal diverse ECM roles in directing cancer cell invasion.

## 3. Discussion

This study represents, to our knowledge, the first application of hybrid scaffolds allowing independent control of ECM biochemical and biophysical properties, decoupling collagen concentration from bulk and microscale mechanical stiffness. While hybrid scaffolds combining hydrogels and porous scaffolds have been developed previously, most examples have used synthetic scaffolds or fibres primarily as mechanical reinforcement, with limited emphasis on cell-matrix interaction.^20,21^ Here, we demonstrate a direct link between hybrid scaffold composition and cell morphology, in systems with constant bulk mechanical properties. Other examples have focussed on mimicking a specific tissue of interest, for instance Offeddu *et al.*^22^ combined poly(vinyl alcohol) (PVA) and poly(acrylic acid) (PAA) with freeze-dried collagen scaffolds to mimic cartilage’s osmotically stiffened structure. The novelty of our current study lies in the range of biochemical and biophysical properties explored, demonstrating a three-fold increase in hybrid scaffold stiffness at constant composition, and a six-fold increase in collagen concentration with only a modest accompanying change in stiffness (from ∼3 to 4 kPa).

Interestingly, our results also demonstrated the crucial role of ECM viscoelasticity in determining cell behaviour. Surprisingly, we found that fibroblasts encapsulated within hybrid scaffolds containing a collagen gel (Hy-CG) did not elongate, unlike their behaviour in both collagen gels and scaffolds alone. Application of particle tracking microrheology to examine biomaterial mechanical properties on the single cell scale revealed that while the collagen gel local shear modulus *G’* remained constant with and without the presence of the ice-templated scaffold, the loss modulus *G’’* and loss tangent tanδ were both significantly lower in the Hy-CG relative to the gel alone (Supplementary Fig.4). This indicates a lower contribution from the viscous component of the Hy-CG mechanical response, i.e. energy dissipation. Previous studies have demonstrated the link between energy dissipation and cell elongation, indicating that viscous response is in many cases necessary for cell elongation in 3D culture.^23,24^ While in this study we chose to focus on hybrid scaffolds containing peptide gels for invasion analysis (Hy-PG), since these did not show the same drop in viscous response, several techniques exist for modulating collagen gel viscoelasticity that could be used for further exploration of the Hy-CG for 3D cell culture, such as control of incubation time before gelation or application of different crosslinking modalities.^3,25^

Application of Hy-PG to investigate breast cancer cell growth and morphology demonstrated that while MDA MB 231 did not elongate in peptide gels alone, as has been previously described,^15^ Hy-PG supported their elongation onto the ice-templated collagen fibres. This was accompanied by a significant increase in cell metabolic activity over time, whereas MDA MB 231 in PG alone showed only a small and non-significant increase. Hy-PG also supported growth and elongation of MCF7, and importantly maintained the characteristic growth morphology of both cell lines, with MCF7 (estrogen/progesterone receptor positive) forming dense colonies and MDA MB 231 (estrogen/progesterone receptor negative) growing as stellate single cells, reflecting their typical morphologies in 3D culture.^18^

Finally, we demonstrated that patient-derived breast cancer cells, sourced from PDX material as described previously,^26^ invade in response to hybrid scaffolds of controlled ECM composition and structure. We have previously shown that peptide gels demonstrate tuneable ECM composition, which may be controlled independently of stiffness.^17^ Here we show an added dimension to these degrees of freedom, incorporating a peptide gel with invasion-promoting composition within a hybrid scaffold, allowing simultaneous control over collagen fibre structure. Control over the solute chosen for ice templating has previously been shown to influence the long-range invasion and cell migration speed of fibroblasts and MC3T3 preosteoblasts, by influencing pore interconnectivity.^19,27,28^ Here, we demonstrate that scaffolds templated using collagen suspended in 0.05M acetic acid guide the directional invasion of patient-derived breast cancer cells, with collective cell invasion following the contact guidance from the patterned collagen fibres. Collective breast cancer invasion has previously been shown to be collagen I dependent, while interactions with other ECM components such as fibronectin are also known to play an important role.^29,30^ The ability of the hybrid scaffold system to independently tune such properties highlights its exciting potential for decoupling diverse ECM roles in breast cancer development, as well as in other applications in health and disease.

## 4. Conclusion

Here we have demonstrated, for the first time, application of hybrid scaffold systems for decoupling biophysical and biochemical ECM roles in determining cell phenotype. Combining defined hydrogels with ice-templated collagen scaffolds, these hybrid scaffolds permit independent control over ECM composition and stiffness, with their macroscopic dimensions maintained over culture through carbodiimide crosslinking. We have demonstrated the suitability of the hybrid scaffolds for probing the correlation between ECM biochemical and biophysical properties and cell phenotype, providing the tuneablity to match these properties to different soft tissues of interest. We suggest that these hybrid scaffolds form a suitable platform technology to allow future investigation of the role of ECM in health and disease, across a range of applications from cancer to wound healing.

## 5. Experimental Section

### Ice-Templated Scaffold Fabrication

Bovine fibrillar collagen type I (Collagen Solutions) was swollen at 1% (w/v) in either 0.001 M hydrochloric acid (HCl) or 0.05 M acetic acid, incubated overnight at 4°C, and homogenised using a benchtop blender. Batch testing was carried out to ensure that each suspension could be pipetted accurately, applying an additional dialysis step for highly viscous suspensions, in which the collagen was dissolved in 0.01 M sodium hydroxide (NaOH) and dialysed against deionised water for 2 days. The freeze-dried final product was then swollen in acid as described above, resulting in lower viscosity and improved pipetting accuracy. Suspensions were briefly centrifuged to remove bubbles, before transferring into 12-well cell culture plates (4 mL/well) or into custom made steel moulds (10 mm filling height).^31^ These were rapidly cooled to -20°C and -35°C respectively to induce freezing, before ice sublimation at 0°C and 80 mTorr for 1200 minutes.

Scaffolds were chemically crosslinked with 1-ethyl-3-(3-dimethylaminopropyl)-carbodiimide hydrochloride (EDC, Sigma-Aldrich, UK) and N-hydroxysuccinimide (NHS, Sigma-Aldrich, UK) in 75-95% (v/v) ethanol.^32^ A molar ratio of 5:2:1 of EDC:NHS:COO-(Collagen) is referred to as 100% cross-linking, decreasing to 50%, 25% and 10% cross-linking at molar ratios of 5:2:2, 5:2:4 and 5:2:10 as previously described.^33^ Scaffolds were immersed in fresh cross-linking solution at room temperature for 2 hr under agitation, followed by at least 4 washes in deionised water to remove excess ethanol. The scaffolds were freeze-dried again following the same protocol as before and stored at room temperature until needed.

### Hydrogel Fabrication

Synthetic peptide hydrogels were fabricated as described previously.^15^ Briefly, 12.5 mg FEFEFKFK peptide was dissolved in 800 µL sterile water, vortexed, and heated to 80°C for 2 hours to induce peptide dissolution (Peptimatrix Ltd., UK). The pH was then adjusted using 0.5 M NaOH and 100 µL 10X phosphate buffered saline (PBS) until optically clear, before incubating at 80°C overnight. The resulting gel precursors were stored at 4°C until needed, heating to 80°C directly before use, and equilibrated at 37°C at the point of seeding. 250 µL cell culture medium (high glucose Dulbecco’s Modified Eagle Medium with 10% FBS and 1% L-glutamine (DMEM) unless specified) was added immediately prior to plating and mixed slowly with a reverse pipetting technique to avoid bubbles. The final peptide concentration was 10 mg/ml. To create the invasive peptide gel composition, extracellular matrix components were diluted in cell culture medium to 500 µg/ml collagen I (07005 Stem Cell Technologies, UK), 50 µg/ml vitronectin (A14700, Fisher, UK) and 50 µg/ml fibronectin III (a kind gift from Professor Clair Baldock).^34^ This was added to the peptide gel precursor at a 1:5 dilution, resulting in final concentrations of 100, 10 and 10 µg/ml in the final gel respectively.

Collagen gels were fabricated by pH adjustment of an acidic collagen solution (Sigma-Aldrich, UK). 6 mg/ml collagen solution was diluted to the desired concentration with 0.01M HCl. The pH was then adjusted using 0.1 M NaOH (15% total volume) and 10X PBS (10% total volume) to give the gel precursor, immediately before plating.

### Hybrid Scaffold Fabrication

Ice-templated scaffolds were cut into 8 mm diameter cylinders with a biopsy punch, and sectioned at 1-4 mm, either manually with dissection scissors, or using a Leica VT1000S Vibratome. Scaffolds were sterilised in 70% ethanol and washed with either PBS or cell culture medium as appropriate for the cell type in culture (DMEM unless specified). Scaffolds were squeezed to remove excess liquid before perfusing with peptide or collagen gel precursor, applying a gentle pressure to the scaffolds while pipetting the gel precursor into the scaffold structure. 250 µL precursor per 4 mm scaffold section was used unless specified. The resulting hybrid scaffolds were incubated at 37°C to induce gelation. Cell culture medium (DMEM unless specified) was added after 15 minutes. Samples containing peptide gels were given a further 2 media changes within the next hour.

### Scaffold Morphological Analysis

Scaffold samples were prepared for Scanning Electron Microscopy imaging by sputter coating with a 5 nm layer of platinum. Imaging took place in secondary electron mode, at an accelerating voltage of 5 kV, using a FEI XL30 system. Images were processed using the OrientationJ plugin in ImageJ ^35^ for visualisation of orientation patterns within the sample.

For quantitative analysis of fluorescent bead position within hybrid scaffolds, samples were prepared incorporating 6 µm diameter fluorescent beads (Polysciences) by mixing into the gel precursors at 3×10^5^/mL. Beads in PBS alone were also added as a control. Samples were imaged using an Ultima2Pplus multiphoton microscope, Bruker, UK, at a laser excitation wavelength of 740 nm. Z-stacks were taken at a step size of 10 µm and distances between beads and the closest pore wall were manually measured using Fiji software.^36^

### Bulk Compression Analysis

Ice-templated scaffolds, hydrogels and hybrid scaffolds were prepared as described above, using scaffolds sectioned at 4 mm. Samples were incubated at 37°C in DMEM overnight before testing. All tests were conducted using a Universal Testing Machine (1ST, Tinius Olsen) with a load cell of 25 N. Sample diameter and thickness were measured with Vernier callipers and a ruler before the test for calculation of applied stress (force/area). Compression tests were performed along the height of the sample at a crosshead speed of 2 mm/min. Stress-strain curves were plotted and the Young’s modulus estimated from the gradient of the initial linear region of the curve, with a minimum of 4 measurements per condition.

### Passive Microrheology

Ice-templated scaffolds used for microrheology were sectioned to 2 mm thickness. Hydrogels and hybrid scaffolds were prepared for microrheology as described above, incorporating 6 µm diameter polystyrene beads (Polysciences) at a final concentration of 1.5 × 10^6^/mL by mixing into the gel precursors. Samples were plated at 200 µL per well of an 8-well glass-bottom IBIDI chamber slide (Thistle Scientific, UK), either alone or as a hybrid scaffold. Samples were incubated at 37°C to induce gelation. Samples were immersed in DMEM with 1% HEPES to maintain hydration and pH, and all testing took place in a humidified, environmentally controlled sample chamber at 5% CO^2^ and 37°C.

Single particle tracking microrheology data was acquired from encapsulated beads using the OptoRheo platform described previously for video microscopy, and custom acquisition software previously developed in Micro-Manager.^37–39^ Each bead was brought into focus to achieve an image with a bright centre, and the bead position across 500,000 frames was found using real-time image processing and recorded to disk for later analysis.. Narrowband noise due to electronic interference was removed from the position-time data using MATLAB’s bandstop function. This was visualised by calculating the power spectral density (PSD) of the position data using the expression *PSD* 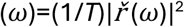, where *T* is the total measurement duration and *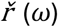* is the Fourier transform of the position-time series (evaluated using the FFT algorithm). A region of width 1 Hz was manually chosen to remove each noise peak that was consistently observed in the PSD across all measurements.

The normalised mean-squared displacement (NMSD, Π(τ)) was calculated at logarithmically spaced lag times τ, according to Π(τ)=⟨(*r*(*t*)−*r*(*t*+τ))^2^⟩^t^ / ⟨*r*^2^⟩_*t*_, where the numerator is the mean-squared displacement (MSD) with position *r*(*t*) at time *t*, position *r*(*t*+τ) at a time point τ seconds later, the angled brackets ⟨ ⟩_*t*_denoting an average over all possible values of time *t*; while the denominator is the MSD value averaged across a range of lag times τ corresponding to the plateau MSD. The resulting MSD data was exported into the MATLAB graphical user interface π-Rheo to calculate plots of *G’* (storage modulus) and *G’’* (loss modulus) for each bead, with a minimum of 8 measurements per condition.^37,40,41^

### Cell Maintenance

Human dermal fibroblasts (HDF, European Collection of Animal Cell Cultures, UK), MCF7 and MDA MB 231 (National Cancer Institute, USA) were cultured in DMEM as described above, with 1% penicillin/streptomycin (P/S) added for HDF culture and for all cell seeding experiments. HDF were used at passage 8-14, and MCF7 and MDA MB 231 were used below passage 30. All cells were cultured at 5% CO^2^ and 37°C in a humidified atmosphere.

Cells extracted from patient derived xenografts (PDX) were also used in this work. PDX lines, derived in-house,^26,42^ were maintained in immunodeficient Rag2-/-γc-/-mice (RAG2G, 8–10 weeks) bred in-house under PPL P375A76F, according to United Kingdom Home Office Animals (Scientific Procedures) Act 1986 recommendations. Mice were housed in individually ventilated cages (Tecniplast, UK) within a barriered unit illuminated by fluorescent lights on a 12-hour cycle (on 07.00, of 19.00), air conditioned by a system set to maintain 21 ± 2 °C and a humidity of 55% ± 10%. Mice were housed in social groups with irradiated bedding, autoclaved nesting materials and environmental enrichment (Datesand, UK). Sterile irradiated 5V5R rodent diet (IPS Ltd, UK) and irradiated water (SLS, UK) were offered ad libitum. Only female mice were used as the tissue being maintained was from breast tumours. Tissue was generated by serial passage of tumour fragments using an implant trochar (VetTech Ltd., UK) into the mammary fat pad of the mice with 50 µl Matrigel™, by licenced competent in vivo technicians under project license PPL 3003444. Tumours were measured weekly using Vernier callipers, and the volumes were calculated using the formula V = ab2/6, where a is the length and b is the width. Mice were weighed weekly and given a daily health check by an experienced technician. All in vivo work was approved by the University of Nottingham AWERB.

Single cells for ex vivo culture were isolated from tumours by mincing with a scalpel and incubation in 12 ml collagenase/dispase solution (collagenase type II 100 U/ml, dispase 2.4 U/ml in HBSS, Invitrogen). After 80 min at 37 °C with rotation, the sample was washed through a 70 µm nylon mesh with phenol red free RPMI (R7509 Sigma), followed by centrifugation (300g, 5 minutes) to extract the cell pellet. Extracted cells were then cryopreserved in 90% FBS with 10% DMSO, or maintained in ex vivo culture using serial passage in peptide hydrogels as previously described until required for seeding.^26^

### Analysis of HDF Morphology

Cells were stained with Invitrogen CellTracker™ (Fisher, UK), to enable visualisation by fluorescent microscopy. Stocks were made up in DMSO according to product instructions, before final dilution to 2.5 µg/ml in DMEM-PR (no phenol red/FBS, with only 1% L-glutamine, 1% P/S and 1% sodium pyruvate, Sigma-Aldrich, UK). A flask of HDF was washed with DMEM-PR and incubated with the staining solution for 30 minutes, before removal from the flask using 1x trypsin-EDTA, and counting to seed at 1×10^5^ cells per sample. Samples were cultured for 3 days before a 2-day fixation in 4% (w/v) paraformaldehyde to allow multiphoton imaging. Samples were washed and stored in PBS prior to imaging.

Samples were imaged using an Ultima2Pplus multiphoton microscope, Bruker, UK, at a laser excitation wavelength of 810 nm. Maximum intensity Z-projections were created from Z-stacks taken from the top, middle and bottom of the scan, each with 30 slices (120 μm). Scans were taken from the surface of the sample to a depth of around 500 µm, with a step size of 4 µm. To reveal the cell distribution across the entire sample (X-Y plane), tile scans were performed using Atlas volume imaging with 5% tile overlap. For quantitative analysis of cell morphology, multiphoton images were processed in Fiji software ^36^ to allow automatic cell detection. Images were sharpened, Otsu thresholded and despeckled to remove noise, and particle detection was set between 30-1000 pixel^2^, and circularity 0.2-1, to exclude background noise and scaffold struts from the cell analysis. For Atlas images, tiles were stitched together using the grid/collection stitching plugin. Linear blending was used for fusion with a regression threshold of 0.3.

### PrestoBlue™ Viability Assay

MCF7 and MDA MB 231 cells were harvested using TrypLE (Fisher, UK) and suspended in peptide gels, prepared as described above, or in DMEM alone, at a concentration of 5×10^5^ cells/mL. The cell-loaded peptide gels were plated at 100 µL/well of a 96-well plate, and also used to make hybrid scaffolds as described above, adding 125 µL gel per scaffold sample. For the scaffold only condition, the cell suspension in DMEM was added at 125 µL per scaffold. Peptide gels and hybrid scaffolds were also prepared using DMEM alone to provide a cell-free control. All samples were incubated at 37°C for 15 minutes to promote cell attachment, before gently adding DMEM to cover the samples. Two additional media changes in the next hour were given to samples containing peptide gels.

The PrestoBlue™ HS Cell Viability assay (Fisher, UK) was used to measure cell metabolic activity at day 0 (3 hours after seeding), 4 and 7. Scaffolds were moved into a fresh plate to avoid interference from any cells adhered to the base of the well. PrestoBlue™ solution was added to each sample at 1:10 dilution and incubated for 1 hour in the dark at 37°C. 100 µL aliquots were taken from each sample in duplicate and plated in a 96-well plate, for fluorescence detection using a FluoStar Omega plate reader, excitation/emission 544/590, gain 1000. Background values as measured for the cell-free control were subtracted from all samples, and measurements were normalised to readings taken for each condition at day 0, to account for any difference in cell seeding eficiency.

### Breast Cancer Cell Line Imaging

Samples seeded with MCF7 or MDA MB 231 as described above were fixed for 1 hour in 10% formalin at day 7 after seeding, and stored in PBS. Samples were incubated for 1 hour in 0.5% bovine serum albumin and 0.1% Triton X100 in PBS, before staining with rhodamine-Phalloidin (1:400, Fisher, UK) and 1:1000 DAPI solution (D3571, Fisher, UK), allowing visualisation using a Leica TCS SPE confocal microscope.

### Spheroid Invasion Analysis

For all PDX experiments, including hybrid scaffold preparation, the medium used was phenol red free RPMI-1640 medium containing 10% FBS and 1% L-glutamine. PDX cells harvested as described above were seeded at 2000 cells per well in an ultra-low attachment plate (Corning 7007), in 200 µL medium/well. Plates were centrifuged at 300 g for 10 minutes to promote spheroid formation. At day 7 after seeding, 100 µL medium was removed per well and replaced with 100 µL CellTracker™ dye in phenol red free RPMI-1640 (with 1% L-glutamine) at 5.5 µg/ml (final concentration 5 µM). Spheroids were incubated for 90 minutes to allow penetration of the dye throughout the spheroid. After staining, the spheroids and staining solution were transferred to a falcon tube using a P1000 pipette tip with the end removed with dissection scissors, and the mixture briefly centrifuged to isolate the spheroids. At least 5 spheroids were added to each hybrid scaffold, prepared as described above. Hanging inserts (662610 Greiner) were found to improve retention of the spheroids on the scaffold surfaces. The seeding surface was imaged using a Nikon Eclipse widefield microscope at regular timepoints in culture. Spheroid size was measured at each time point by tracing the spheroid perimeter from the darkfield images, using the wand tool in Fiji software as previously described.^42^ Orientation coherency was measured for each spheroid from brightfield images, using the OrientationJ - Measure plugin.^43^ Three experimental repeats were carried out using PDX cells derived from three separate patients.

### Statistical Analysis

Statistical analysis took place using GraphPad Prism version 10.4.1. An unpaired t-test or one-way ANOVA with a Tukey test for multiple comparisons were used as appropriate, with the assumption of normality verified using the Shapiro-Wilk test. Statistical significance was declared at p<0.05.

## Acknowledgements

This work was funded by the University of Nottingham (Anne McLaren fellowship, JCA), Nottingham Breast Cancer Research Centre (JCA, AMG, CLRM), and St John’s College, Cambridge (Benefactors’ scholarship, XS). The Henry Royce Institute for advanced materials Equipment Access Scheme supported access to the Cambridge 3D Bioelectronics Facility, through Cambridge Royce facilities grant EP/P024947/1 and Sir Henry Royce Institute recurrent grant EP/R00661X/1. The authors thank the University of Nottingham Nanoscale and Microscale Research Centre (nmRC) for providing access to instrumentation, and Ms Nicola J. Weston for technical assistance.

