## Supplementary data for "Hybrid Scaffolds Decouple Biochemical & Biophysical Regulation of Cell Phenotype"

**Co-corresponding authors:*

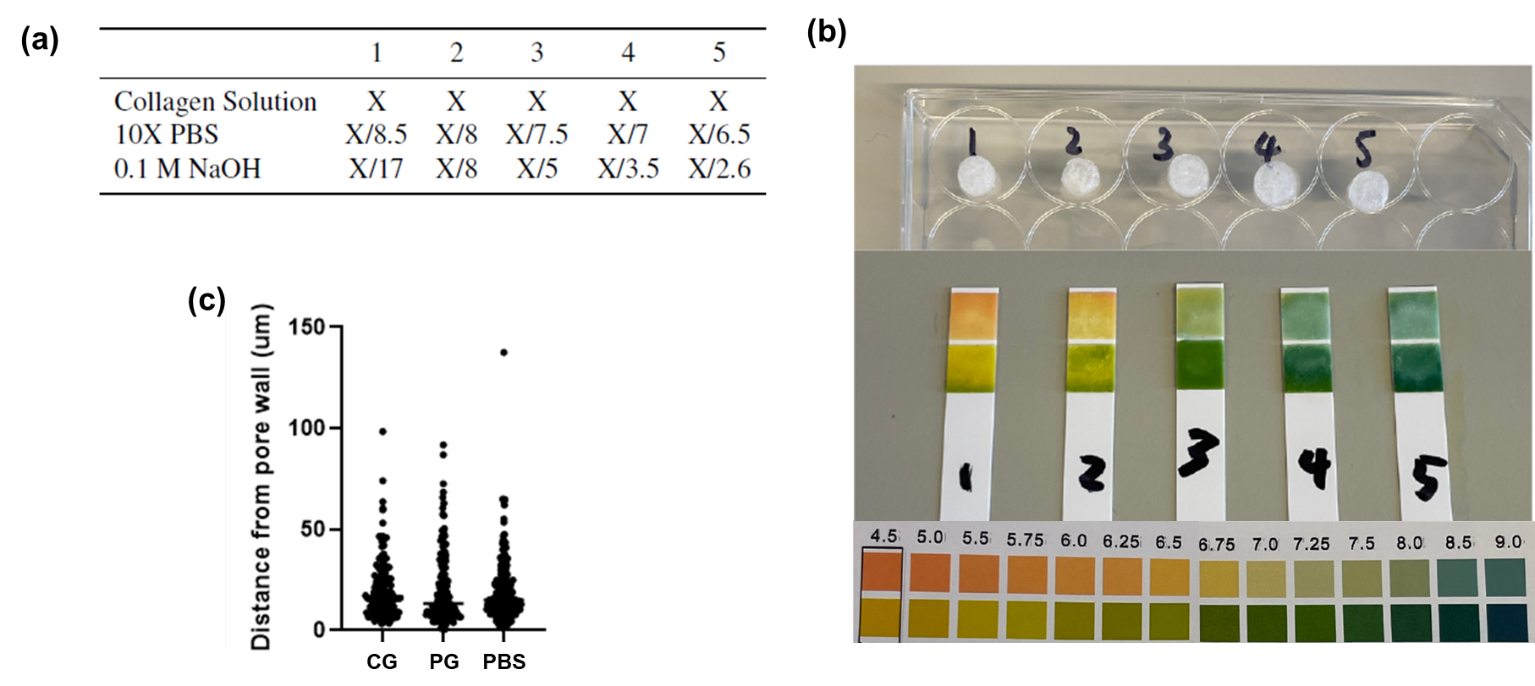

**Supplementary Figure 1: Optimisation and validation of hybrid scaffolds.** (a) Table showing collagen gel formulations investigated for optimum gelation within hybrid scaffold; (b) pH measurements demonstrating the need for additional NaOH for collagen neutralisation in hybrid scaffolds relative to published protocols (formulation 1); (c) Quantification of bead distribution within hybrid scaffolds shows that both collagen gels (CG) and peptide gels (PG) penetrate effectively into the pore structure, showing no difference in bead positioning to a bead suspension in PBS alone.

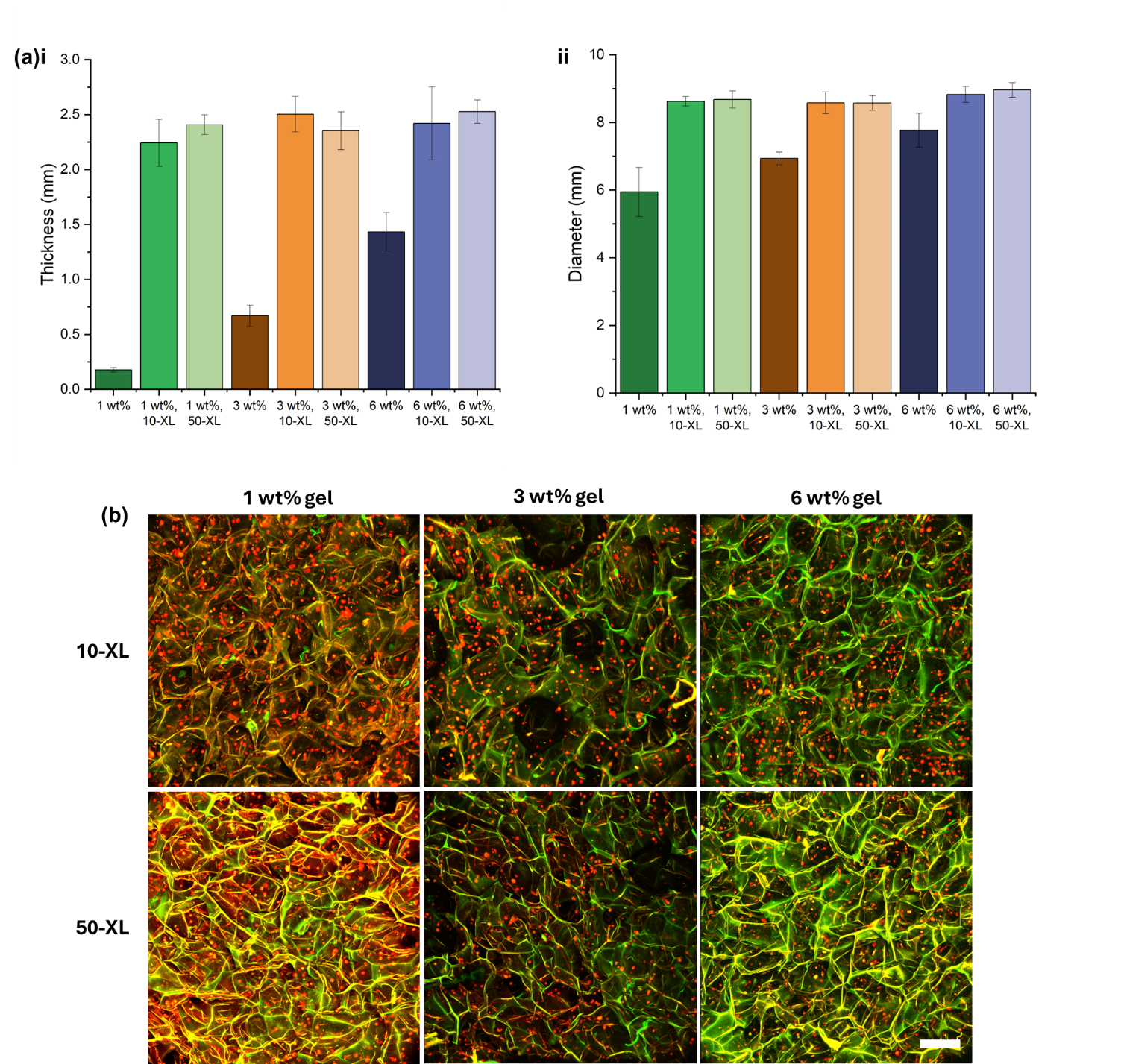

**Supplementary figure 2:** Hybrid scaffolds containing collagen gels (Hy-CG) show no contraction over culture with human dermal fibroblasts (HDF). (a) Measurements of (i) thickness and (ii) diameter of CG and Hy-CG with varying collagen concentration and scaffold crosslinking (XL, numbers indicate percentage crosslinking) reveal that contraction only occurs in CG alone, and not in Hy-CG. (b) Multiphoton images show that HDF cultured within Hy-CG have a rounded morphology across all collagen concentrations and crosslinking levels tested. Scale bar 200 µm.

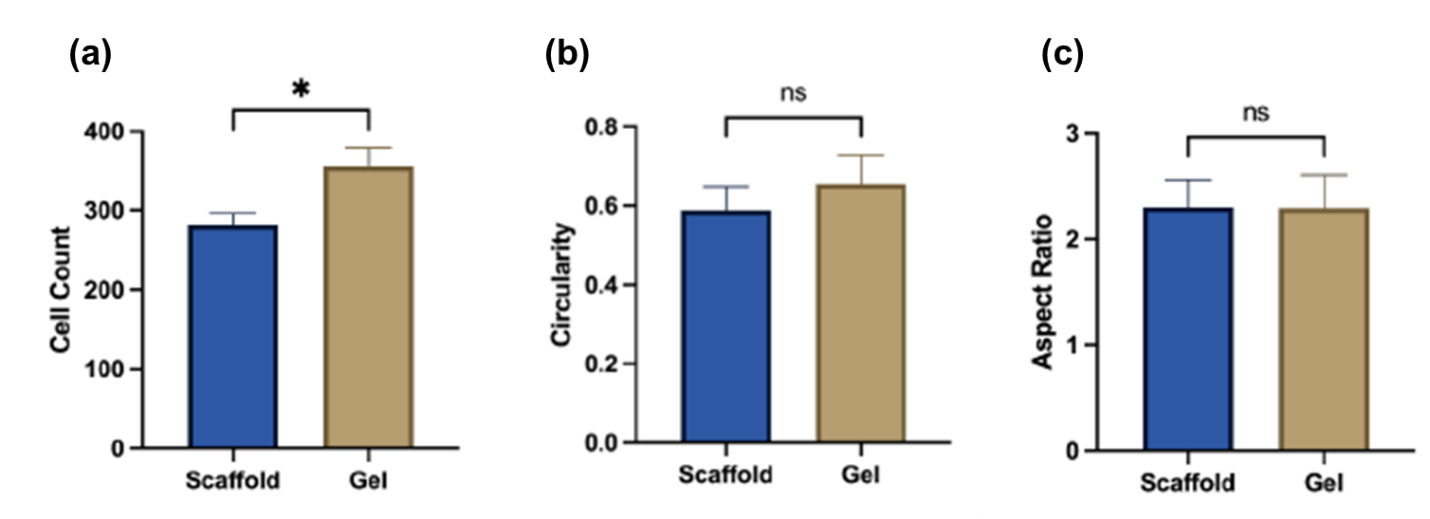

**Supplementary Figure 3:** Cell seeding method does not affect human dermal fibroblast (HDF) morphology. (a-c) Comparison of cell count, circularity and aspect ratio of HDFs seeded into Hy-CG via the scaffold or the CG component. Cells were cultured for 2 days.

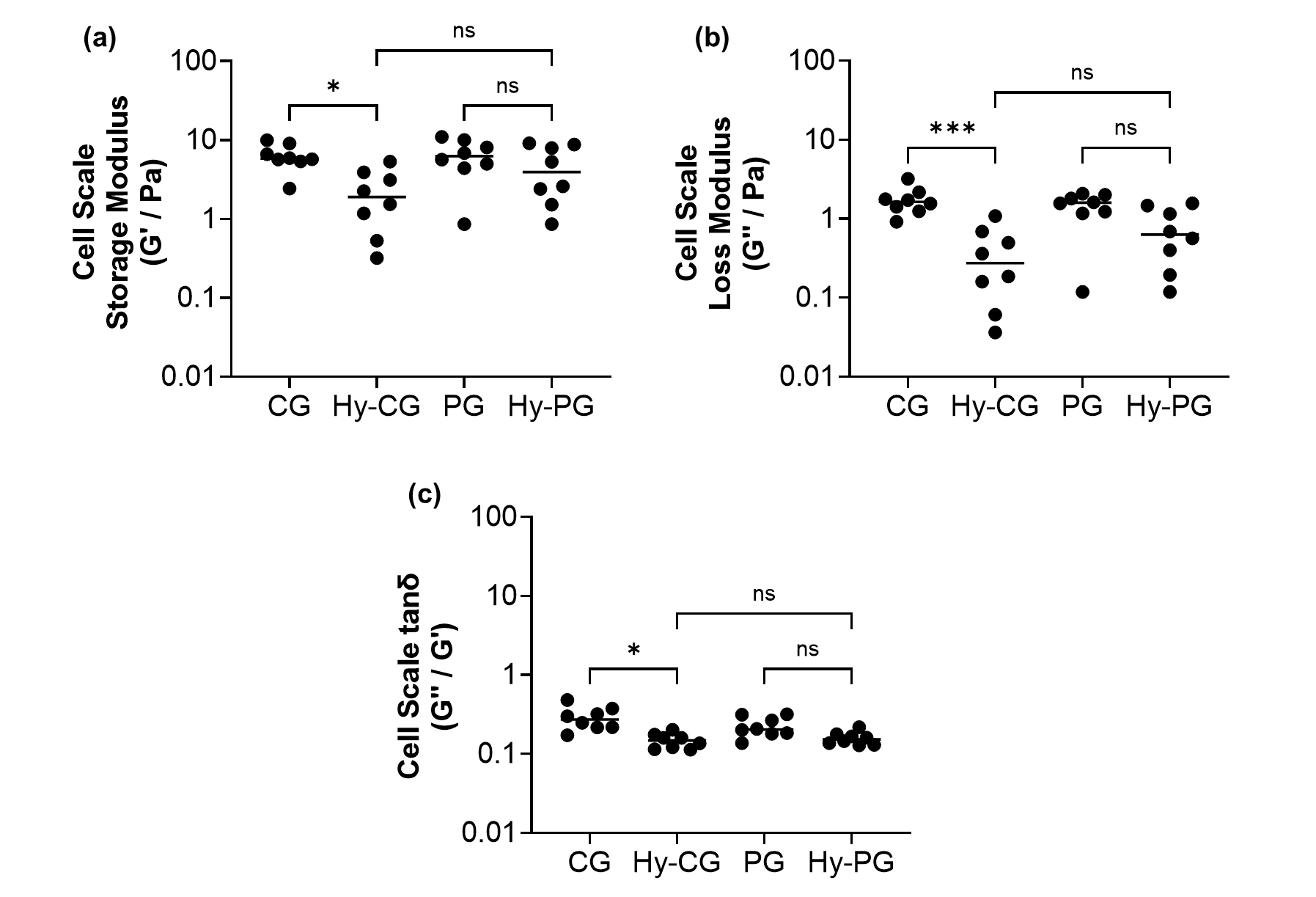

**Supplementary Figure 4: Local mechanical response differs between CG and Hy-CG but not between PG and Hy-PG**. (a) Measurements of storage modulus G’, (b) Loss modulus G’’ and (c) Loss tangent tanδ reveals no significant difference in any of these measurements between PG and the corresponding hybrid scaffold Hy-PG, in contrast to the comparison between CG and Hy-CG. Measurements for 8 beads from two samples per condition are shown.
